# Single-cell RNA sequencing of conditional *Pten* knockout neurons in mice

**DOI:** 10.1101/2022.03.17.484809

**Authors:** Marek Svoboda, Bryan Luikart, Giovanni Bosco

**Affiliations:** Quantitative Biomedical Sciences Program, Geisel School of Medicine, Dartmouth College; Molecular and Systems Biology Program, Geisel School of Medicine, Dartmouth College

**Keywords:** single-cell RNA sequencing, NGS, PTEN, Conditional Knockout, mouse

## Abstract

PTEN is a well-known tumor suppressor whose mutations are also strongly associated with Autism Spectrum Disorder (ASD). The activity and function of PTEN in neurons have been studied extensively in various settings, whereas animal models offer the best opportunity to experimentally modify and test the role of PTEN in neuronal development *in vivo*. On the molecular level, PTEN’s importance in the mTOR pathway’s suppression is well-known but many other interactions have been suggested and yet others likely remain undiscovered. Therefore, to systematically explore the regulatory landscape downstream of PTEN on the genetic level, we have set out to establish a singlecell RNA sequencing (scRNA-seq) workflow to measure gene expression changes as a result of knocking out the *Pten* gene *in vivo*. We were able to collect brain tissue from four conditionally knocked out mouse samples and dissociate them into single live cells, which were subsequently flow-sorted and a scRNA-seq library was prepared. All of these steps were completed in a single day, yielding sequencing data from hundreds of cells suitable for cell clustering, cell type identification, and differential expression measurements. Having demonstrated its feasibility, this approach promises to aid hypothesis generation and discovery in mapping the function of *Pten* as well other genes whose regulatory networks remain to be fully defined.

## Introduction

Phosphatase and TENsin homolog (*PTEN*) was first identified as the gene responsible for Cowden syndrome located on chromosome 10q22-23 (1). Its function was soon established as a tumor suppressor missing in endometrial carcinoma (2), glioma (3), kidney (4), breast (5), and prostate cancers (6, 7). Dominant germline *PTEN* mutations have been also linked to other tumor syndromes: Bannayan-Riley-Ruvalcaba syndrome (8), Proteus syndrome (9), and Proteus-like syndrome (10). Subsequently, these phenotypically heterogeneous diseases were combined under the diagnostic umbrella of *PTEN* Hamartoma Tumor Syndrome (PHTS) (11, 12). Soon after its discovery, mutations in *PTEN* were also associated with autism spectrum disorder (ASD) (13). *PTEN* mutations are now identified in about 10% of people with ASD and macrocephaly (14, 15), making it one of the most commonly mutated genes in these patients, which account for about 20% of all ASD cases (16). Apart from other pathogenic signs like seizures, ataxia, and Lhermitte-Duclos disease (17), macrocephaly is one of the hallmark signs of individuals carrying germline *PTEN* mutations, with or without PHTS, underscoring the importance of this gene in neurons, where it has been found to be important for normal function, development, and morphology (18–20) and its loss leads to their precocious differentiation, mislocation, hypertrophy, hyperexcitability, and increased survival (21–26).

Several mouse models have been created to study PTEN function. While homozygous *Pten* loss leads to embryonic lethality, *Pten* heterozygous mice have high tumor incidence (27–29). Mouse models for studying Pten’s role in neurons include its conditional knockout (cKO, Pten^flox/flox^) (30), short hairpin RNA knockdown (24), and nuclear excluded PTEN (Pten^M3M4^) (31). These mice reproduce the phenotypes seen in PTEN-ASD patients including macrocephaly, seizures, and abnormal social interactions (16, 23, 32).

Thanks to these mouse models, we can study PTEN function on the cellular and molecular level. PTEN has been described as a lipid and protein phosphatase (17, 33). Its most studied function is dephosphorylation of PIP_3_, which results in regulation of the PI3K-Akt-mTORC1 pathway (34, 35) (Figure 1A). Accordingly, many of the *Pten* KO-associated changes are rescued or prevented when neurons are treated with rapamycin, an mTOR inhibitor (36). However, previous work shows that excessive microtubular (MT) polymerization, which mediates abnormal neural process formation in *Pten* KO neurons, is not rescued by acute rapamycin treatment (Figure 1B, 37). This suggests that PTEN exerts its regulatory activity in neurons through other pathways.

**Figure 1.**
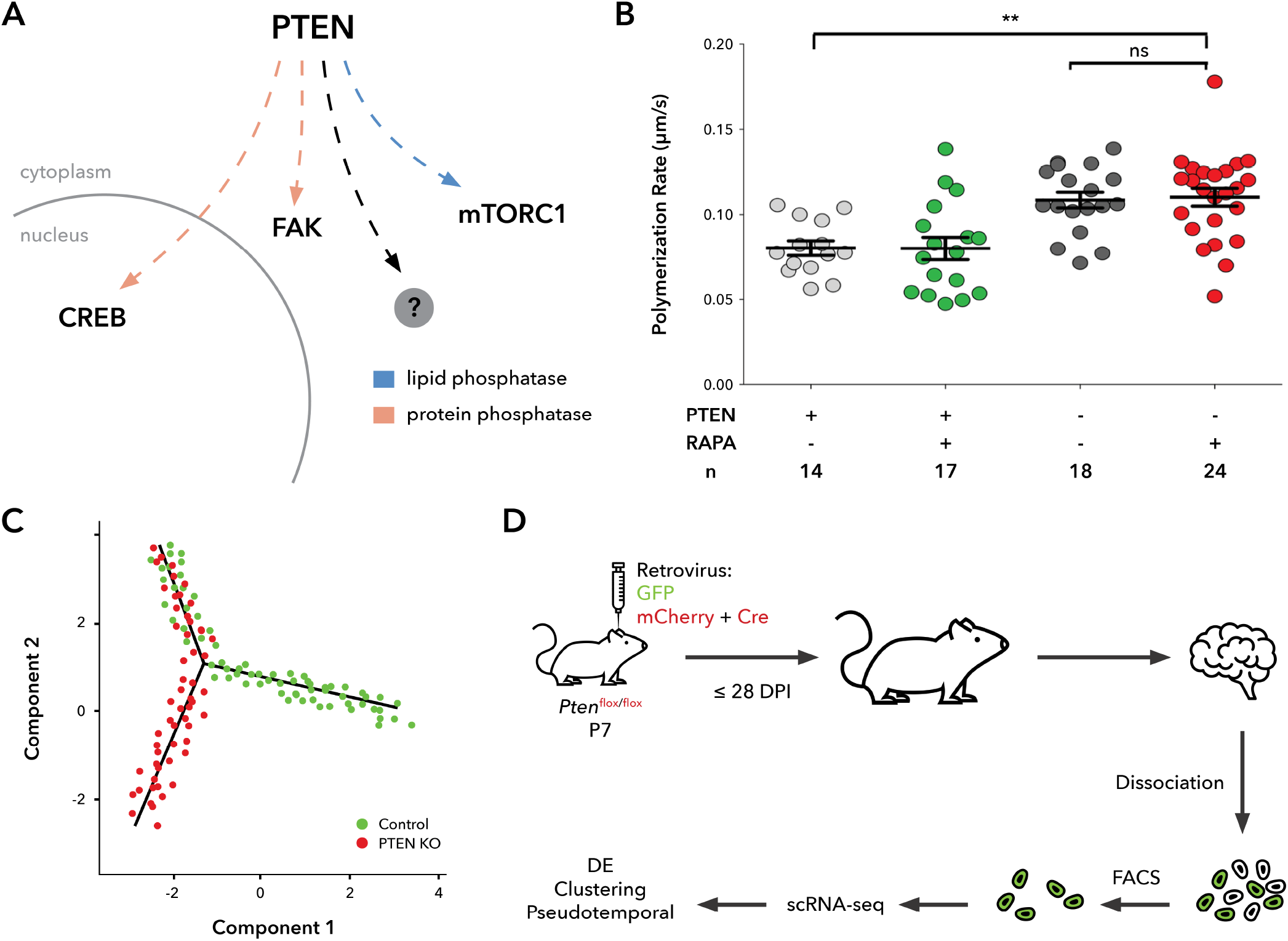
Identification of novel *Pten*-dependent pathways using scRNA-seq. **A.** Neuronal pathways regulated by PTEN lipid and protein phosphatase activity. While the effects of PTEN’s lipid phosphatase activity on the mTORC1 pathway have been well studied, little is known about the significance of PTEN’s protein phosphatase activity. It has been suggested to regulate cell migration through focal adhesion kinase (FAK)8 and neural stem cell differentiation through cyclic AMP response element-binding (CREB) protein (40). **B.** Acute rapamycin treatment (“RAPA +”) does not correct the microtubular polymerization rate in *Pten* KO (“PTEN -”) neurons. Graphical representation of data published by Getz et al. (37). **C.** The expected outcome of pseudotemporal ordering of individual cells using the Monocle 2 (44) algorithm. *Pten* KO neurons are expected to deviate from Control neurons’ developmental trajectory of ordered cell states. **D.** Flowchart outlining the experimental design: mice are injected with retroviral particles containing either pRubiG (GFP) or pRubiC-T2A-Cre (mCherry and Cre separated by a self-cleaving peptide, T2A) around post-natal day 7 (P7). Cells are collected 28 days post-injection (28 DPI), dissociated, and enriched for GFP+ (control) or mCherry+ (KO) cells before scRNA-seq and subsequent computational analysis.

While the effects of PTEN’s *lipid* phosphatase activity on PIP_3_ have been well studied, little is known about the significance of PTEN’s *protein* phosphatase activity. PTEN protein phosphatase has been suggested to regulate cell migration through focal adhesion kinase (FAK) (38) and Shc (39) in glioblastoma (GBM), neural stem cell differentiation through cyclic AMP response element-binding (CREB) protein (26, 40) (Figure 1A), neuronal spine density (41) and tumor suppression of GBM (42) by autodephosphorylation, and cell metabolism through insulin receptor substrate-1 (IRS-1) (43).

Additionally, interactions with other genes seem to be of great importance for the penetrance of PTEN-associated pathology. For example, males are four times more likely than females to be diagnosed with ASD (45, 46) and similar pattern is found in PTEN-specific cases (13, 47, 48). In mice, behavioral deficits in both *Pten* haploinsufficient (32) and cytoplasm-predominant (49) models are sex-specific as well. Similarly, penetrance of *Pten* haploinsufficiency-associated tumors depends on genetic background (50). Nuclear localization of PTEN (51) and its interactions with multiple targets (38, 40) indicate it may have indirect effects on transcriptional regulation. Taken together, these studies strongly suggest important interactions between *PTEN* and other genes crucial for ASD and other pathologies, whose molecular mechanisms are yet to be elucidated.

Despite the abundant evidence that PTEN interacts with other proteins and genes, our knowledge of PTEN’s causal involvement in neural disease phenotypes is lacking. Only a systematic investigation of PTEN’s regulatory effects accounting for their diversity and temporal dynamics can ensure that we obtain a complete picture of its downstream mediators and hence potential pharmacological targets. Therefore, we developed a single-cell RNA sequencing (scRNA-seq) protocol to measure gene expression in a conditional knockout (cKO) *Pten* mouse model of autism to study transcriptional states during neuronal development *in vivo* (52). This approach will allow us to analyze the resulting data using a variety of computational methods, such as differential expression (DE), cell clustering, cell type assignment, and pseudotemporal ordering (Figure 1C), to identify the cell states present in *PTEN*-associated ASD and the regulatory pathways downstream of PTEN through which they arise. To our knowledge, scRNA-seq has not been previously used to study transcriptomic changes in selectively *Pten*-disrupted cells.

## Results

### Infection and collection of *Pten* conditional knockout mouse neurons developed *in vivo*

We used homozygous *Pten* conditional knockout (cKO) mice, in which Pten gene exons 4 and 5 were flanked by loxP sequences (Pten^flox/flox^, Figure 1D) (30). This portion of *Pten*, encoding the core phosphatase motif, is excised in the presence of Cre, resulting in *Pten* KO. We targeted hippocampal dentate gyrus (DG), which is one of the structures impaired in ASD (53) and whose continued neurogenesis into adulthood (in both mice and humans) facilitates study of neuronal development and makes it a potential candidate for future treatment targeting. DG in these mice was therefore injected on post-natal day 7 with retroviral particles, which selectively target dividing cells. These particles carried either Cre and mCherry (KO) or only GFP (control). This strategy allows for precise control over the cell population and timing of the KO. Brain tissue was subsequently collected 28 days post injection, eclipsing the peak neurogenesis in mouse hippocampus (54) and allowing any morphological changes to take place after *Pten* KO (25).

The collected brain tissue was subsequently separated into single cells through gentle dissociation. We optimized the timing of this process through several iterations to be able to process up to four samples in a single day to minimize any batch effects and at the same time, to maximize the number of surviving cells for each sample. The following samples were collected: a GFP+ male control (M_GFP), a Cre-mCherry+ male KO (M_mCherry-Cre), a GFP+ female control (F_GFP), and a Cre-mCherry+ female KO (F_mCherry-Cre). All samples had to be staggered while being processed during each precisely timed individual step to ensure that all the steps starting with tissue collection of *in vivo* neurons up to single-cell RNA library preparation could be carried out in a single day. Keeping the timing at a minimum aimed to limit the effect of handling on the final measurement of gene expression in the live neurons.

### Cell sorting through flow cytometry

The dissociated cells were then subjected to fluorescence-activated cell sorting (FACS) using the appropriate lasers and gating parameters to select only those single cells (i.e. singlets) that survived the extraction and dissociation process described above (i.e. low DAPI) and had been fluorescently labeled *in vivo* (GFP or mCherry positive for the control or KO cells, respectively).

Aiming to extract as many live neurons as possible, we were able to dissociate about 1,000,000 cells from each of the four samples studied. Cells in each sample were then sorted based on four sequential gates in the following combinations of parameters, which were consistent across all samples: FSC (forward scatter) -A (area) vs. SSC (side scatter) -A, which yielded between 11–15.6% of all cells; FSC-H (height) vs. FSC-W (width), which yielded 96.7–99.6% of the remaining cells; SSC-H vs. SSC-W, which yielded 95.6–98.5% of the remaining cells; and GFP (green fluorescent protein) -A vs. DAPI-A, through which only 0.8–1.7% of the remaining cells were selected (Table 1 and Figure 2). This resulted in only 0.1–0.3% of the total number of cells having been positively selected by FACS, yielding approximately between 1,000–3,000 of the original about 1,000,000 total cells sorted per sample.

**Table 1.** Proportions of cells filtered by FACS in sample 2F_GFP. Table showing the numbers and relative (%Parent) and absolute (%Total) proportions of cells selected by each subsequent FACS gate in a sub-sample of 10,000 cells, as shown in Figure 2.

| Population | #Cells | %Parent | %Total |
| --- | --- | --- | --- |
| All Events | 10,000 | NA | 100.0 |
| P1 (FSC-A vs. SSC-A) | 1,559 | 15.6 | 15.6 |
| P2 (FSC-H vs. FSC-W) | 1,541 | 98.8 | 15.4 |
| P3 (SSC-H vs. SSC-W) | 1,513 | 98.2 | 15.1 |
| P4 (GFP-A vs. DAPI-A) | 26 | 1.7 | 0.3 |

**Figure 2.**
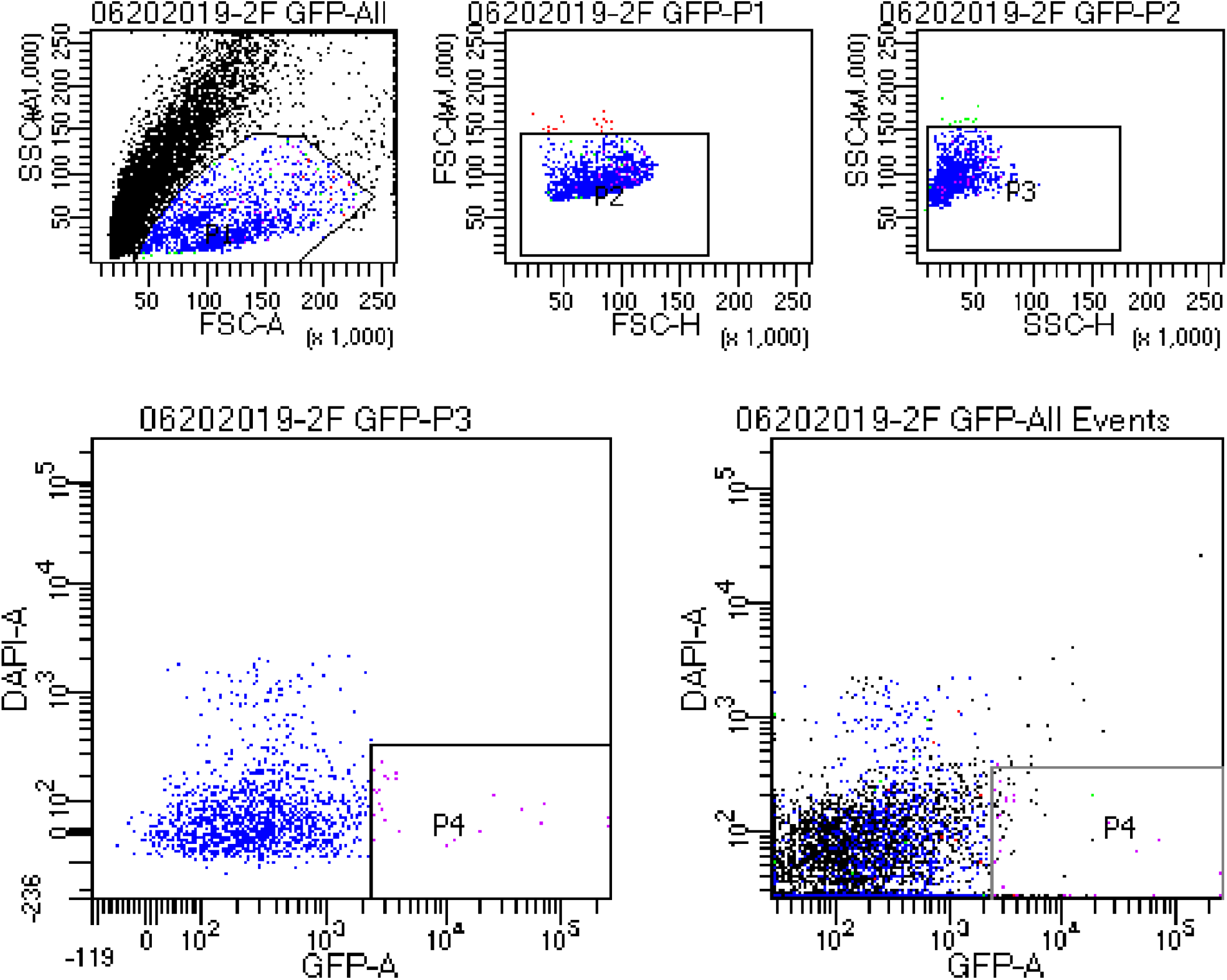
Fluorescence-activated cell sorting (FACS) of cKO neurons from sample 2F_GFP, using four separate gates. The four gates used for cell sorting. Top: P1 - FSC (forward scatter) -A (area) vs. SSC (side scatter) -A, singlets inside gate; P2 - FSC-H (height) vs. FSC-W (width), singlets inside gate; P3 - SSC-H vs. SSC-W, singlets inside gate. Bottom: P4 - GFP (green fluorescent protein) -A vs. DAPI (4’,6-diamidino-2-phenylindole) -A, GFP positive alive cells inside gate; left: only cells selected by all previous gates are shown, right: all cells are shown.

### Single cell RNA sequencing quality control filtering

The flow-sorted cells were then used for single-cell RNA library preparation using the standard 10X Genomics Chromium protocol followed by next-generation sequencing. Sequencing reads were subsequently decomplexed and aligned to the mouse genome, yielding a gene-cell matrix for each sample. The final number of cells detected by single-cell RNA sequencing (scRNA-seq) per sample ranged between 314–640, the mean number of sequencing reads per cell was between 143,802–329,980, and the median number of genes detected per cell was between 27–2,198 (Table 2). Based on the significantly lower median numbers of genes detected per cell, the samples from the female mice were determined to be of insufficient quality and were not included in the subsequent analyses.

**Table 2.** Summary statistics for the scRNA-seq samples before filtering. The columns denote the mouse sex, sample condition, total number of cells, the mean number of reads per cell, and the median number of genes per cell detected in each respective scRNA-seq library.

| Sex | Condition | Cells | Reads/c | Genes/c |
| --- | --- | --- | --- | --- |
| M | GFP (Control) | 314 | 307,449 | 1,378 |
| M | mCherry-Cre (KO) | 322 | 329,980 | 2,198 |
| F | GFP (Control) | 640 | 143,802 | 27 |
| F | mCherry-Cre (KO) | 377 | 299,249 | 95 |

As is customary in scRNA-seq data analysis (55), the cells were further filtered based on the metrics discussed above, as well as the content of mitochondrial genes detected (Figure 3A). Because too few genes per cell are usually associated with low cell quality or background measurement and too many genes per cell suggest the presence of doublets, we removed cells that contained less than 200 or more than 5,000 genes (Figure 3B). Similarly, cells with an excessively high proportion of mitochondrial genes suggest loss of mitochondrial integrity associated with cell death and therefore contamination secondary to cell membrane disruption. Hence, cells that contained more than 20% mitochondrial genes were also removed. After this quality control (QC) filtering step, 193 control and 240 KO cells from the male samples were left in the dataset.

**Figure 3.**
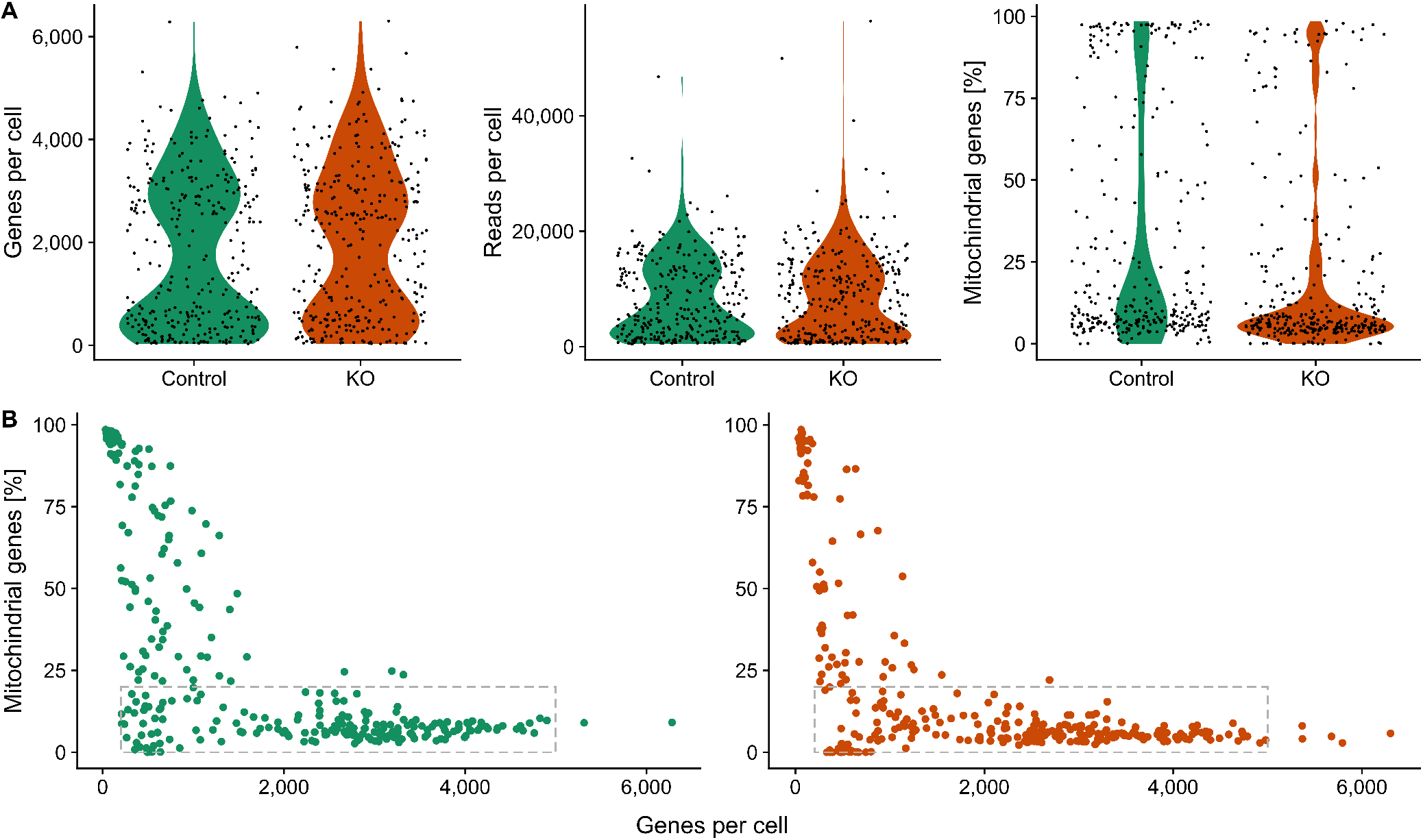
Quality control (QC) measurements of the male samples. **A.** Violin plots of the number of genes, total number of RNA reads, and percentage of mitochondrial genes per cell in each sample. Samples depicted: 1M_GFP (control - green) and 5M_mCherry (KO - orange). **B.** Plots of the number of genes vs. percentage of mitochondrial genes per cell in each of the samples. Cells enclosed in the gray boxes are those that passed the QC filtering - i.e., containing between 200–5,000 genes and less than 20% of mitochondrial genes detected. The colors are the same as in **A.** - left: control; right: KO.

**Figure 4.**
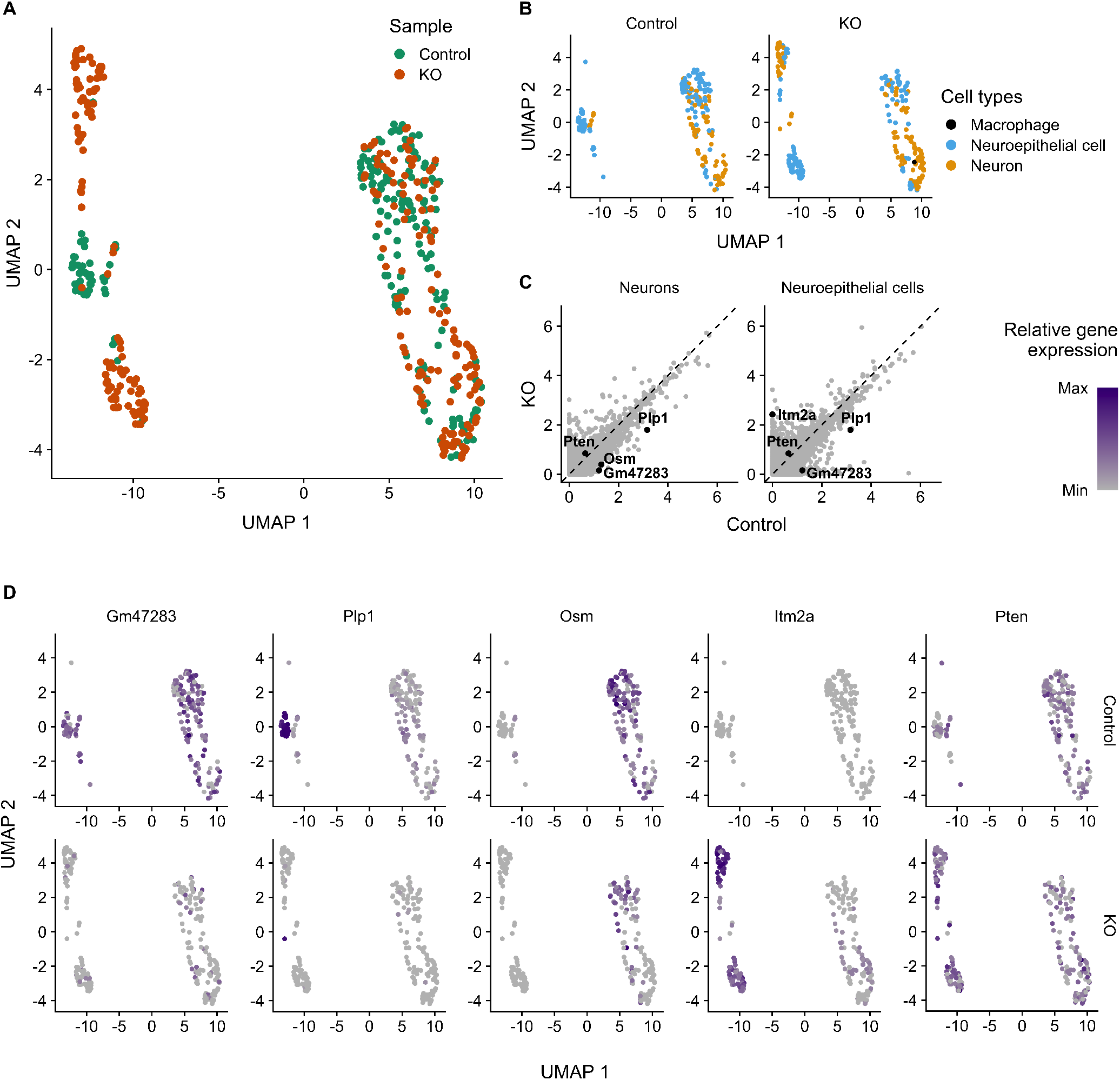
Clustering analysis of the male samples. **A.** UMAP plot of all the cells that passed the QC, colored by the sample or origin. **B.** UMAP plots of cells split by sample and colored by the cell types detected: neurons, neuroepithelial cells, and a macrophage. (Only one cell in the KO sample and none in the control sample was classified as a macrophage.) **C.** Plots of normalized gene expressions in the control vs. KO sample, in the two major cell types detected. *Pten* and the top three genes most significantly differentially expressed between the samples in neurons and neuroepithelial cells are highlighted. The gray dashed line represents equal expression between the samples. **D.** UMAP plots colored by the relative expression of the genes highlighted in **C.**, split by the sample of origin. The differential expressions between the control vs. KO sample of all these genes were statistically significant, except for *Pten*.

### Cell clustering and gene expression patterns detected

Clustering analysis of the *Pten* KO (n = 240) and control (n = 193) neurons from the male mice showed significant overlap between *Pten* KO and control cell populations, suggesting that transcriptome of some cells did not change significantly after *Pten* KO (Figure 3A). Notably, however, each of the two experimental groups also exhibited a transcriptionally unique cell population not found in the other group, suggesting a detectable effect associated with knocking out of *Pten* in a subset of cells. Namely, the control sample contained a cluster of neuroepithelial cells not found in the KO sample and, similarly, the KO sample contained one cluster of neuroepithelial cells and one cluster of neurons not found in the control sample (Figure 3B). Knockout of *Pten* during neuronal development may therefore lead to a change in the cell differentiation process, whereby the transcriptional state of some neuroepithelial cells (i.e., the neural stem cells) is altered and may also lead to the development of neurons with an abnormal transcriptional profile.

*Gm47283* and *Plp1* were the top two most significantly differentially expressed genes between the control and KO samples in both neurons and neuroepithelial cells, followed by *Osm* and *Itm2a* in these cell types, respectively (Figure 3C). The *Plp1* gene, which is X-linked and encodes for the proteolipid protein integral to myelin formation (56), was most highly expressed in the neuronal cluster specific to the control sample (Figure 3D). Conversely, the *Itm2a* gene, which encodes for Integral membrane protein 2A shown to be important in chondro-osteogenic and myogenic differentiation (57), was most highly expressed in the two clusters specific to the KO sample. It is also notable that the gene that was one of the most differentially expressed genes in neurons but not in neuroepithelial cells, *Osm*, encodes the Oncostatin M protein, which has been found to suppress the growth of several oncogenic cell lines (58), activity comparable to that of the tumor suppressor PTEN. The *Pten* gene was not detected as differentially expressed between the control and KO samples in any of the cell types detected, nor among all the cells overall.

## Discussion

As the role of PTEN in neurons remains to be elucidated on the molecular level, in this study, we have developed an unbiased approach to explore the regulatory pathways downstream of PTEN in *vivo* and demonstrated the feasibility to collect up to four samples in a single day using this protocol. Specifically, we explored the KO-induced cellular changes using scRNA-seq to capture the alterations in transcriptional states during neuronal development. To our knowledge, *Pten* mutations in neurons have not been previously studied using scRNA-seq.

We have identified an absence of a specific subset of neuroepithelial cells and the emergence of a different group of neuroepithelial cells as well as neurons as a result of knocking out *Pten*. We have also detected genes that were differentially expressed between the samples, some of which were specific to neurons. This data underscores the importance of the scRNA-seq approach, as these differences were only found in a subset of cells and would likely be blunted or completely lost using bulk RNA-seq technology.

Additional scRNA-seq experiments will be necessary to optimize this technique and obtain data from enough cells that pass the downstream quality control (QC) checks for further analyses. One of the potential issues of this study is that *Pten* was not differentially expressed between the samples, as would be expected from its targeted knockout. While the introduction of Cre has been previously shown to significantly decrease the amount of PTEN protein in the cells from the mouse model used (25), it is conceivable that the amount of *Pten* mRNA transcripts is not significantly altered by the excision of the gene’s exons 4 and 5, which would explain the lack of its differential expression on the transcriptomic level. It is possible, however, that instead, the cells analyzed by scRNA-seq in the KO sample did not exhibit the intended suppression of PTEN expression, which might have occurred in several ways.

One of the most significant problems encountered during the tissue preparation for scRNA-seq was the noted faintness of the expressed fluorophores (GFP and mCherry). It is known that the endogenously expressed fluorophores, secondary to the *in vivo* retroviral infection of neurons, are relatively lowly luminescent and are therefore often amplified by immunohistochemistry in the in *vitro* microscopy experiments in our laboratory. During FACS sorting, a bimodal distribution of luminescence would be expected, clearly distinguishing the distinctly luminescent cells from those that do not contain the fluorophore, which was not observed during our experiment’s FACS sorting (Figure 2, gate P4).

Additionally, we were unable to detect Cre in the KO sample as an internal control of a successful viral construct infection and expression. This was most likely because the viral construct used lacks a poly(A) signal, the addition of which would enable the detection of Cre in the sequencing data from future experiments.

We also suspect that the low cell yields between cell dissociation and scRNA-seq preparation were caused by excessive cell death. Cell dissociation of live neurons, which possess numerous processes, is notoriously more problematic than that of regularly shaped cells or those that are less tightly interconnected with their milieu. It is therefore common to extract neuronal nuclei instead, which has several advantages (59). Neuronal dissociation is rendered faster, as it does not require excessive care to maintain the cell membrane integrity to keep cells alive and the nuclear membrane itself is more sturdy. Additionally, nuclei can be readily dissociated from frozen tissue, dispensing with the necessity to synchronize the tissue collection for all samples destined for the same scRNA-seq batch into a single day. In combination with a histone-associated eGFP mouse line, where the fluorophore is highly concentrated in the cell nuclei (60), snRNA-seq is therefore likely the best next step in the future iterations of this approach. Additionally, using the same fluorophore (i.e., eGFP) in both the control and KO cells would further standardize the cell sorting process. The improvements outlined above have the potential to further optimize this innovative approach to therapeutic target discovery by mapping transcriptional pathways associated with a complex neural disease, applicable to the study of genes beyond *Pten*.

## Methods

### Mouse *Pten* cKO and tissue collection

Dentate gyrus (DG) of Pten^flox/flox^ mice was injected with retroviral particles carrying either GFP or mCherry-Cre constructs on post-natal day 7 as described by Williams e? *al*. (25). The tissue was then collected 28 days post-injection, as described by Tasic et al. (61). Briefly, individual mice were anesthetized with 2% tribromoethanol (Avertin), perfused with artificial cerebrospinal fluid (ACSF), decapitated, and the brain was immediately removed and submerged in fresh ice-cold ACSF bubbled with carbogen gas (95% O_2_ and 5% CO_2_). The brain was sectioned on vibratome on ice, and each slice (300 μm) was immediately transferred to an ACSF bath at room temperature. DG was then microdissected under a microscope, and the slices after the dissection were visualized under a confocal microscope to verify the presence of the respective fluorophores. Subsequently, the cell dissociation protocol was carried out using the Papain Dissociation System (Worthington, LK003150), following the manufacturer’s protocol. The dissociation protocol was optimized to yield at least 80% viable cells, as determined using Trypan Blue.

### Flow cytometry and single-cell RNA sequencing

Single cells were isolated by fluorescence-activated cell sorting (FACS) on FACSAria III cell sorter using a 70 μm nozzle, sorting at 70 psi, in single-cell sorting mode, into phosphate-buffered saline (PBS) and stored on ice. To exclude dead cells, DAPI was added to the single-cell suspension to a final concentration of 2 ng/ml and FACS populations were chosen to select single cells with low DAPI (using the 405 nm laser) and high GFP (488 nm laser) or mCherry (561 nm laser) signal. The accuracy of sorting was confirmed by visually observing DAPI staining in the sorted cells. Sorted cells were then processed by the 10X Genomics Chromium workflow for scRNA-seq library construction and subsequently sequenced on Illumina NextSeq 500 sequencer.

### Data analysis

The sequencing files were decomplexed and reads were aligned to the mm10 v3.0.0 Cell Ranger mouse reference genome using Cell Ranger v3.1 software. Subsequent data analysis was carried out in R using the Seurat package v4.0.5 (62). Only features detected in at least three cells were included as were only cells containing between 200–5, 000 genes total and no more than 20% mitochondrial genes. The data was subsequently normalized using regularized negative binomial regression (63). Cell types were detected with the python package ScNym (64), using an adversarial neural network model trained on TabulaMuris brain tissue data (65).

## ACKNOWLEDGEMENTS

This research was supported by Bakala Foundation (M.S.), Rosaline Borison Memorial Fund (M.S.), Burroughs Wellcome Fund - Big Data in the Life Sciences Training Program at Dartmouth (M.S.), National Institutes of Health: DP1MH110234 (G.B.), Geisel School of Medicine at Dartmouth (G.B.). This article was written using the HenriquesLab bioRxiv template.

